# Colibactin-induced DNA damage activates cGAS-STING signaling that is amplified by mitochondrial DNA release

**DOI:** 10.64898/2026.11.25.769242

**Authors:** Shuangshuang Xie, Jasmine Tzeng, Terry D. Kim, Yougant Airan, Seth B. Herzon, Daniel R. Semlow

## Abstract

The microbiome-derived genotoxin colibactin induces oncogenic mutations that contribute to the development of colorectal cancer. Colibactin exposure also elicits inflammatory and senescence-associated responses. However, the cellular pathways that link colibactin-induced genotoxic stress to inflammation and senescence are poorly defined. Here, we show that colibactin causes cytoplasmic DNA accumulation and activates pro-inflammatory cGAS-STING signaling in human cells. Genetic or pharmacological inhibition of cGAS reduces colibactin-induced inflammatory gene expression and the accumulation of senescence- associated markers. We further show that these responses occur in two phases: an early response that is independent of mitochondrial DNA (mtDNA) and a stronger late response that is amplified by mtDNA release. Inactivation of either cGAS or the mitochondrial outer membrane permeabilization factor BAK suppresses cytoplasmic mtDNA accumulation and dampens late inflammatory signaling. Together, our data support a model in which colibactin-induced nuclear DNA damage initially activates cGAS-STING, which promotes BAK-dependent mtDNA release and thereby enhances inflammatory signaling. This work reveals a mechanism through which a clinically relevant microbial genotoxin reprograms host cell physiology and may contribute to the development or progression of colorectal cancer.

**Importance Statement:** Colibactin is a DNA-damaging agent produced by certain bacteria implicated in colorectal cancer and other diseases. Exposure to colibactin causes cancer-associated mutations and has also been linked to inflammatory responses. However, the mechanisms that connect colibactin-induced DNA damage to inflammatory signaling remain unclear. Here, we report that colibactin promotes inflammation by activating the cGAS-STING pathway, an innate immune signaling pathway that senses cytoplasmic DNA. This response is further enhanced by the programmed release of mitochondrial DNA into the cytoplasm. These findings identify a mechanism by which colibactin-induced DNA damage triggers and amplifies inflammatory signaling, potentially contributing to colorectal cancer development.

## Introduction

Host-microbiome interactions are important determinants of human health and disease. Accumulating evidence indicates that genotoxic agents produced by gut bacteria contribute to pathological states, including inflammatory bowel disease (IBD) and colorectal cancer (CRC) (1). Among the best-studied microbiome-derived genotoxins is colibactin, a secondary metabolite produced by *Escherichia coli* (*E. coli*) and other *Enterobacteriaceae* that harbor the *pks* polyketide synthase–nonribosomal peptide synthetase biosynthetic gene cluster (2). Colibactin-producing *pks+* bacteria are detected in approximately 66% of biopsies from CRC patients, and colibactin-associated mutational signatures are detected in approximately 5% of human CRC tumors (3, 4). Colonization with *pks+ E. coli* promotes intestinal tumorigenesis in mice (3), and chronic exposure to *pks+ E. coli* induces CRC driver mutations in human colonic organoids (4). Even transient exposure to colibactin is sufficient to induce genomic instability in primary colonic cells (5). Taken together, these findings implicate colibactin-induced DNA damage as an important contributor to CRC development.

Colibactin consists of a labile central scaffold with two cyclopropane groups, each of which can alkylate DNA (6–8). Colibactin preferentially reacts with the N3 positions of offset deoxyadenosines on opposite DNA strands to form highly toxic DNA interstrand cross-links (ICLs) that block DNA replication and transcription (2). Additionally, colibactin can monoalkylate DNA and promote depurination, resulting in a spectrum of DNA lesions (9). Consistent with this mechanism of action, colibactin exposure produces characteristic T>N single-base substitutions and single-nucleotide deletions at AT-rich sequence motifs. Although colibactin-induced ICLs encountered during replication can be repaired by the Fanconi anemia ICL repair pathway, colibactin exposure nevertheless causes DNA double-strand breaks, G2/M cell-cycle arrest, and mitotic abnormalities (10, 11). These findings help explain how colibactin can cause host cell genome instability and generate CRC-associated driver mutations.

In addition to destabilizing the host cell genome, colibactin has been linked to cellular inflammation and senescence, both of which can promote carcinogenesis (12, 13). In mouse models of CRC and colitis, colonization with *pks+ E. coli* increases histological evidence of inflammation, including immune-cell infiltration (14, 15). Exposure to *pks+ E. coli* or a synthetic colibactin analog also induces the expression of pro-inflammatory cytokines (3, 16). Moreover, human cells cocultured with *pks+ E. coli* acquire features of cellular senescence, including decreased proliferative capacity, accumulation of p21 and phosphorylated p53, and increased senescence-associated β-galactosidase (SA-β-gal) activity (17, 18). Colibactin-induced inflammation and senescence may therefore provide additional mechanisms by which *pks+* bacteria contribute to the pathogenesis of CRC and other disease states.

Although colibactin exposure has been associated with increased inflammation and senescence, the mechanisms linking colibactin to these phenotypes are poorly defined. In general, genotoxic stress can activate inflammatory signaling through multiple pathways, among which the cGAS-STING pathway is one of the best-characterized (19–21). Cyclic GMP-AMP synthase (cGAS) is a pattern-recognition receptor that binds double-stranded (ds)DNA in the cytoplasm. DNA binding activates cGAS to synthesize 2’,3’-cyclic GMP-AMP (cGAMP), a second messenger that binds and activates the endoplasmic reticulum (ER)-anchored adaptor protein stimulator of interferon genes (STING). cGAMP binding induces STING oligomerization and recruitment of TANK-binding kinase 1 (TBK1). TBK1 phosphorylates the transcription factor IRF3, which translocates to the nucleus and activates expression of type I interferons and interferon-stimulated genes. STING also activates NF-κB signaling through IκB kinase (IKK), thereby promoting expression of pro-inflammatory cytokines. Although the cGAS-STING pathway was originally characterized as an innate immune defense that responds to pathogen-derived DNA, it can also be activated by self-DNA that gains access to the cytoplasm. During genotoxic stress, such DNA can arise through rupture of micronuclei, mitochondrial damage, or release of nuclear chromatin. Persistent DNA damage can also trigger inflammatory signaling through cytoplasmic RNA-sensing pathways or through mechanisms independent of cytoplasmic nucleic acid accumulation (22–25). Whether any of these pathways contribute to colibactin-induced inflammation and senescence remains unclear.

Here, we investigated how colibactin-induced DNA damage promotes inflammatory signaling in human cells. We show that both a synthetic colibactin analog and *pks+ E. coli* induce cytoplasmic DNA accumulation, activate the cGAS-STING pathway, and promote inflammatory and senescence-associated responses. We further find that cGAS and the mitochondrial outer membrane permeabilization factor BAK promote colibactin-induced release of mitochondrial DNA (mtDNA), thereby enhancing a late inflammatory response. Overall, this study defines a signaling circuit that links colibactin-induced genotoxic stress to inflammation and senescence.

## Results

### Colibactin activates pro-inflammatory signaling through the cGAS-STING pathway

To determine whether colibactin contributes to inflammatory signaling, we first examined induction of type I interferon and pro-inflammatory cytokine genes in HT-29 human colon cancer cells treated with the synthetic colibactin analog clb742. HT-29 cells were selected for this analysis because they retain functional STING-dependent signaling, which is defective in many other colon cancer cell lines (26). clb742 contains an unsubstituted ethylene spacer in place of the labile diketone moiety found in native colibactin and is otherwise identical to the natural product (Fig. 1A) (27). Like coculture with *pks+ E. coli*, clb742 induces cell cycle arrest in human cells and T>N mutations in mice (16, 27). Moreover, we have shown that ICLs generated by native colibactin and clb742 are both repaired by the Fanconi anemia ICL repair pathway (28). These findings support the use of clb742 to model colibactin-induced changes in host cell physiology.

**Fig. 1.**
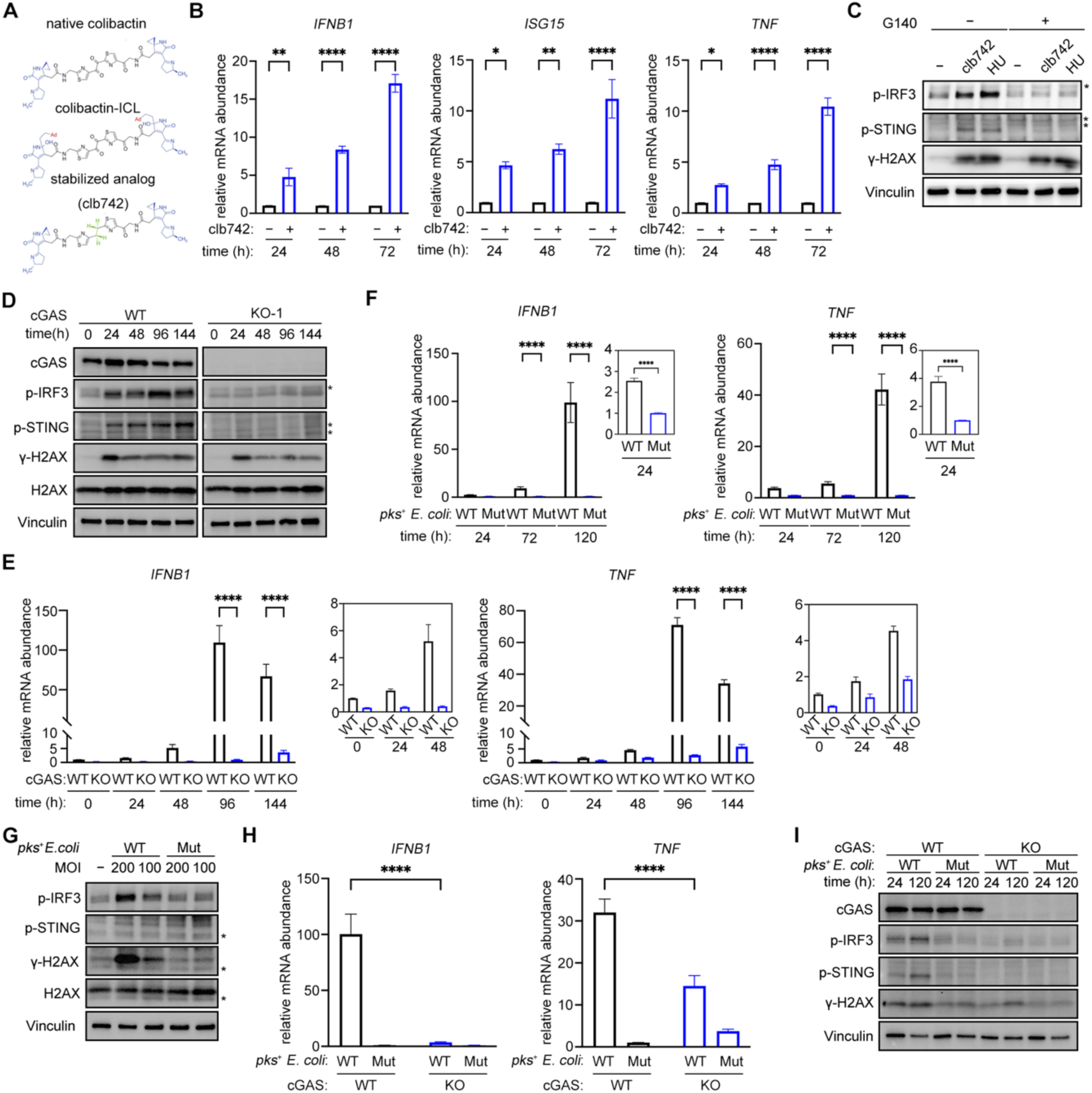
Colibactin activates the cGAS–STING pathway. **(A)** Chemical structures of colibactin, a colibactin-induced DNA interstrand cross-link (ICL), and the synthetic clb742 colibactin analog. The position of the two-carbon spacer that replaces the α-aminoketone present in native colibactin is indicated in green. Ad, adenine. **(B)** HT-29 cells were treated with clb742 for the indicated times and *IFNB1*, *ISG15*, and *TNF* mRNA expression was determined by quantitative RT-PCR. Relative mRNA expression at each time was normalized to the untreated condition. Results are presented as the mean ± the standard error of the mean (SEM) from 3 independent experiments. Statistical significance was determined using two-way ANOVA followed by Sidak’s multiple-comparisons test, *P* < 0.05 (*)*, P <* 0.01 (**), *P* < 0.001 (***), and *P* < 0.0001 (****). **(C)** HT-29 cells were treated with clb742, HU, and G140 for 24 h, as indicated, and cell lysates were resolved by SDS-PAGE and immunoblotted for the phosphorylated (p-)IRF3, phosphorylated (p-)STING, γH2AX, and vinculin. Asterisks indicate nonspecific bands. **(D)** Wild-type and cGAS-knockout clone 1 (KO-1) HT-29 cells were treated with clb742 for 24 h and then allowed to recover in clb742-free media. At the indicated times following clb742 addition, cell lysates were immunoblotted for cGAS, p-IRF3, p-STING, γH2AX, H2AX, and vinculin as in **C**. **(E)** Wild-type and cGAS-knockout clone 1 (KO-1) HT-29 cells were treated with clb742 for 24 h as in **D** and *IFNB1* and *TNF* mRNA expression was determined by quantitative RT-PCR. Relative mRNA expression was normalized to the untreated (0 h) wild-type condition. Results are presented as the mean ± SEM from 3 independent experiments. Statistical significance was determined as in **B**. **(F)** HT-29 cells were cocultured with wild-type *pks⁺* (WT) or colibactin-deficient *clbP^S95R^* (Mut) *E. coli* at a multiplicity of infection (MOI) of 100 for 4 h and then allowed to recover in *E. coli*-free media. At the indicated times following infection, *IFNB1* and *TNF* mRNA expression was determined by quantitative RT-PCR. Relative mRNA expression was normalized to the 24 h colibactin-deficient *E. coli* condition. Results are presented as the mean ± SEM from 3 independent experiments. Statistical significance was determined as in **B**. **(G)** HT-29 cells were cocultured with wild-type *pks⁺* (WT) or colibactin-deficient (Mut) *E. coli* for 4 h at the indicated MOIs. 24 h after infection, cell lysates were immunoblotted for p-IRF3, p-STING, γH2AX, H2AX, and vinculin as in **C**. **(H)** Wild-type and cGAS-knockout HT-29 single-cell clone (KO-1) were cocultured with wild-type *pks⁺* (WT) or colibactin-deficient *clbP^S95R^* (Mut) *E. coli* at a multiplicity of infection (MOI) of 100 for 4 h and then allowed to recover in *E. coli*-free media. At 120 h following infection, *IFNB1* and *TNF* mRNA expression was determined by quantitative RT-PCR. Relative mRNA expression was normalized to the colibactin-deficient *E. coli* infected wild-type condition. Data are presented as mean ± SEM from 3 independent experiments. Statistical significance was determined as in **B**. **(I)** Wild-type and cGAS-knockout HT-29 single-cell clone (KO-1) were cocultured with wild-type *pks⁺* (WT) or colibactin-deficient *clbP^S95R^* (Mut) *E. coli* at a multiplicity of infection (MOI) of 100 for 4 h and then allowed to recover in *E. coli*-free media for the indicated times. Cell lysates were immunoblotted for cGAS, p-IRF3, p-STING, γH2AX, and vinculin as in **C.**

HT-29 cells were treated with clb742 for up to 72 hours, and reverse transcription quantitative PCR (RT-qPCR) was used to measure relative mRNA abundance of *IFNB1*, a type I interferon gene, *ISG15*, an interferon-stimulated gene, and *TNF*, a pro-inflammatory cytokine gene. clb742 treatment caused a robust, time-dependent increase in expression of all three genes (Fig. 1B). Similar results were obtained with clb742-treated HeLa cells (Fig. S1A), indicating that this response is not specific to HT-29 cells. We conclude that clb742 triggers pro-inflammatory signaling in cultured human cells.

clb742-dependent induction of cytokine genes has been reported to coincide with increased *Cgas* expression in mice, suggesting that cGAS may contribute to colibactin-induced inflammatory signaling (16). We therefore determined whether clb742 activates the cGAS-STING pathway. In both HT-29 and HeLa cells, treatment with clb742 for 24 hours increased phosphorylation of IRF3 and STING, two hallmarks of cGAS-STING pathway activation (Fig. 1C and Fig. S1B). Importantly, clb742-induced IRF3 and STING phosphorylation was suppressed by cotreatment with the cGAS inhibitor G140 and abolished in cGAS-knockout cells (Fig. S1C-E). In contrast, cGAS inactivation did not affect clb742-induced levels of γH2AX, a canonical DNA damage marker, implying that cGAS does not substantially alter clb742-induced nuclear DNA damage accumulation. Depletion of MAVS, the downstream adaptor of the cytosolic RNA sensors RIG-I and MDA5, did not affect clb742-induced IRF3 phosphorylation (Fig. S1F), implying that MAVS-dependent RNA sensing is dispensable for this response. Combined, these data demonstrate that cGAS is required for activation of the STING-IRF3 signaling axis in response to clb742.

We next tested whether cGAS is required for induction of inflammatory genes following clb742 exposure. HT-29 cells were treated with clb742 for 24 hours and then allowed to recover for up to 120 hours in drug-free media. Under these conditions, clb742 induced IRF3 and STING phosphorylation that persisted throughout the recovery period (Fig. 1D). This sustained phosphorylation was inhibited in cGAS-knockout cells. IRF3 and STING phosphorylation was accompanied by a dramatic induction of *IFNB1* and *TNF* expression at 72 and 120 hours after clb742 removal (Fig. 1E). Strikingly, the induction of both genes was attenuated in cGAS-deficient cells. These results establish that the cGAS-STING pathway is an important mediator of the sustained inflammatory response to clb742.

Although clb742 recapitulates many of the genotoxic effects associated with native colibactin, we also tested whether the cGAS-STING pathway is activated in cells exposed to colibactin-producing bacteria. HT-29 and HeLa cells were cocultured for 4 hours with either colibactin-producing *pks+ E. coli* or an isogenic *E. coli* control strain harboring a *clbp^S95R^*mutation that blocks the maturation of active colibactin (29–31). The media was then replaced, and cells were allowed to recover for up to 120 hours. Relative to coculture with the colibactin-deficient control strain, coculture with *pks+ E. coli* induced *IFNB1* and *TNF* expression that continued to increase for 120 hours after exposure (Fig. 1F and Fig. S1G), indicating that bioactive colibactin activates inflammatory signaling in cultured human cells. Coculture with *pks+ E. coli*, but not the control strain, also caused accumulation of γH2AX, phosphorylated IRF3, and phosphorylated STING (Fig. 1G and Fig. S1H). At 120 hours after coculture, cGAS deficiency strongly attenuated *IFNB1* induction and reduced STING and IRF3 phosphorylation (Fig. 1H and I). *TNF* expression was also reduced at this later time point, although the effect was weaker than for *IFNB1*. By contrast, at 24 hours after coculture, *TNF* induction was unaffected by loss of cGAS (Fig. S1I). Thus, sustained inflammatory signaling strongly depends on the cGAS–STING pathway.

### The cGAS-STING pathway promotes colibactin-induced cellular senescence

Next, we asked whether the cGAS-STING pathway contributes to senescence after colibactin exposure. HT-29 cells were once again cocultured with *pks+ E. coli* for 4 hours and then allowed to recover for 120 hours. Senescence was assessed by staining for senescence-associated β-galactosidase (SA-β-gal) activity. Consistent with previous reports (17, 18), coculture with *pks+ E. coli* markedly increased the fraction of SA-β-gal-positive cells relative to coculture with the colibactin-deficient control strain (30% vs 1.8%; Fig. 2A and B). Coculture with *pks+ E. coli*, but not the negative control strain, also resulted in accumulation of p21 (Fig. 2C), a cyclin-dependent kinase inhibitor commonly associated with cellular senescence. Thus, exposure to colibactin-producing bacteria promotes senescence-associated phenotypes in cultured HT-29 cells.

**Fig. 2.**
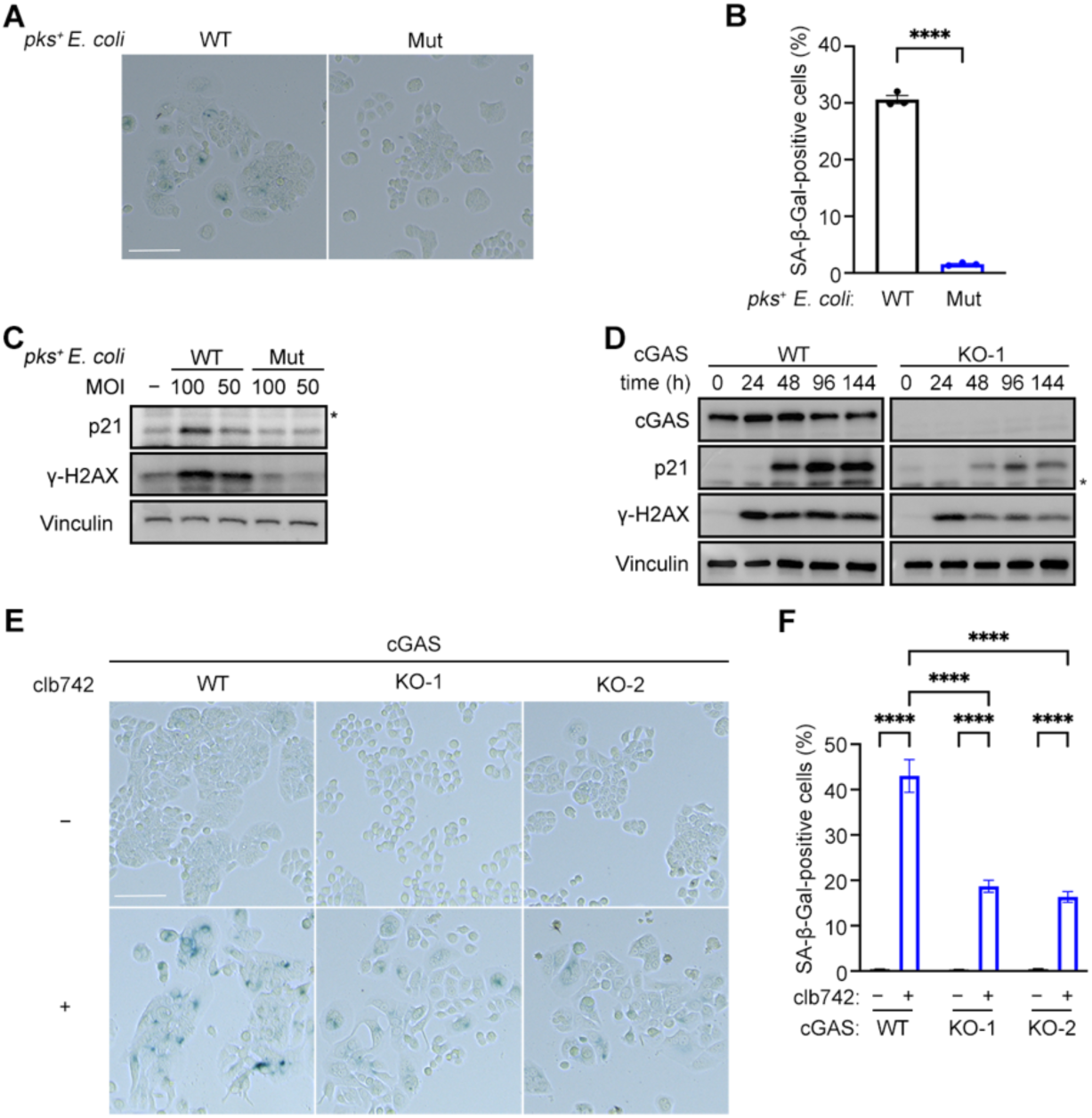
cGAS contributes to colibactin-induced cellular senescence. **(A)** HT-29 cells were cocultured with wild-type *pks⁺* (WT) or colibactin-deficient *clbP^S95R^* (Mut) *E. coli* at an MOI of 100 for 4 h and then allowed to recover in *E. coli*-free media. 120 h after infection, cells were stained for SA-β-gal activity. Representative images are shown. Scale bars, 100 µm. **(B)** Quantification of HT-29 SA-β-gal-positive cells from the experiments described in **A**. Results are presented as the mean ± SEM from 3 independent experiments. Statistical significance was determined using an unpaired two-tailed Student’s *t*-test. *P* < 0.0001 (****). **(C)** HT-29 cells were cocultured with wild-type *pks⁺* (WT) or colibactin-deficient (Mut) *E. coli* for 4 h at the indicated MOIs and then allowed to recover in *E. coli*-free media. 72h after infection, cell lysates were immunoblotted for p21, γH2AX, and vinculin as in Fig. 1C. Asterisks indicate non-specific bands. **(D)** Wild-type and cGAS-knockout HT-29 single-cell clone (KO-1) were treated with clb742 for 24 h and then allowed to recover in clb742-free media. At the indicated times following clb742 addition, cell lysates were immunoblotted for cGAS, p21, γH2AX, and vinculin as in Fig. 1C. **(E)** Wild-type and cGAS-knockout single-cell clone (KO-1 and KO-2) HT-29 cells were treated with clb742 for 24 h and then allowed to recover in clb742-free media. 120 h after clb742 addition, cells were stained for SA-β-gal activity. Representative images are shown. Scale bars, 100 µm. **(F)** Quantification of HT-29 SA-β-gal-positive cells from the experiments described in **E**. Results are presented as the mean ± SEM from 3 independent experiments. Statistical significance was determined using two-way ANOVA with Sidak’s multiple-comparisons test. *P* < 0.0001 (****).

We then tested whether senescence induced by clb742 depends on the cGAS-STING pathway. Similar to coculture with *pks+ E. coli*, clb742 induced p21 accumulation and rendered more than 40% of wild-type HT-29 cells SA-β-gal-positive 120 hours after clb742 exposure (Fig. 2D and F). By contrast, clb742-induced p21 accumulation was diminished in cGAS-knockout cells and the fraction of SA-β-gal-positive cGAS-knockout cells was approximately twofold lower than in wild-type cells (Fig. 2E and F). Notably, despite abolishing clb742-induced phosphorylation of IRF3 and STING, cGAS deficiency did not completely suppress the accumulation of SA-β-gal-positive HT-29 cells following clb742 treatment, implying that clb742 can induce senescence-associated phenotypes through cGAS-independent mechanisms. Nevertheless, these results establish the cGAS-STING pathway as an important mediator of colibactin-induced senescence-associated phenotypes.

### Colibactin exposure triggers cytoplasmic accumulation of nuclear and mitochondrial DNA

Our finding that the DNA-sensing cGAS-STING pathway promotes colibactin-induced inflammation and senescence predicts that colibactin exposure should increase the abundance of cytoplasmic DNA. To test this prediction, we treated HT-29 cells with clb742 for 24 hours and then stained the cells with the dsDNA-specific dye Pico488. Quantification of cytoplasmic Pico488 fluorescence normalized to nuclear fluorescence revealed an approximately twofold increase in signal in clb742-treated cells (Fig. 3A and B), consistent with clb742-induced accumulation of cytoplasmic DNA. A similar increase in cytoplasmic Pico488 intensity was observed in clb742-treated HeLa cells (Fig. S2A and B). In line with a previous report (11), clb742 treatment also increased the fraction of HT-29 cells containing micronuclei (Fig. 3C and D), chromatin-containing extranuclear structures whose membranes are prone to rupture (32) and may contribute to cGAS-STING activation (33–35). clb742 therefore induces both cytoplasmic DNA accumulation and micronucleus formation, either of which could contribute to activation of the cGAS-STING pathway.

**Fig. 3.**
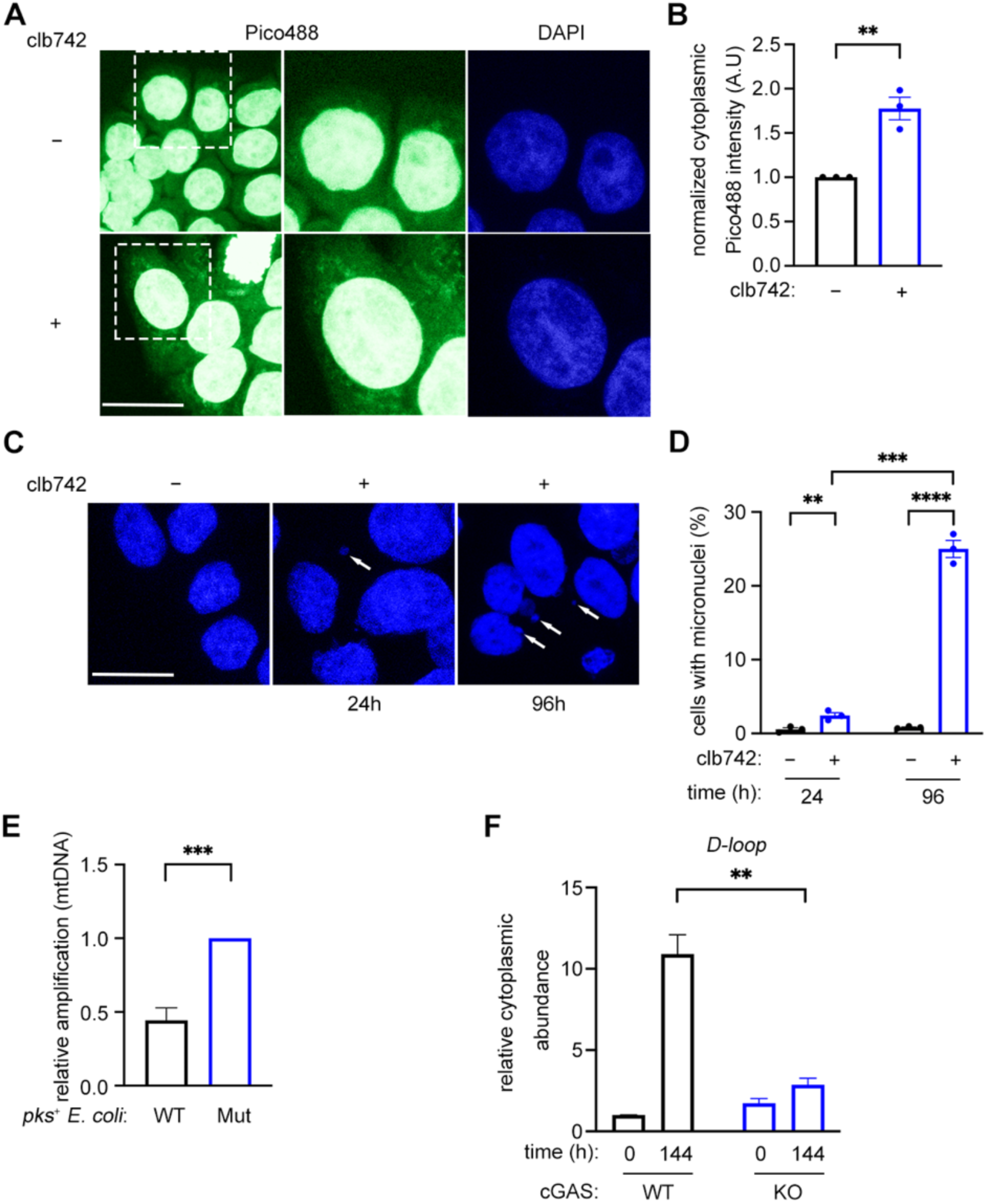
Colibactin induces cytoplasmic DNA accumulation. **(A)** HT-29 cells were treated with clb742 for 24 h and then stained with Pico488 and DAPI. Regions indicated by boxes are enlarged in the middle and right panels. Representative images are shown. Scale bars, 20 µm. **(B)** Quantification of cytoplasmic Pico488 fluorescence intensity from experiments described in **A**. For each cell, cytoplasmic Pico488 fluorescence intensity was normalized to nuclear Pico488 fluorescence intensity. A total of 177–236 cells were analyzed for the untreated group and 185–258 cells for the clb742-treated group in each independent experiment. Results are presented as the mean ± SEM from 3 independent experiments. Statistical significance was determined using an unpaired two-tailed Student’s *t*-test. *P* < 0.01 (**). **(C)** HT-29 cells were treated with clb742 for 24 h or 96 h and then stained with DAPI. Representative images are shown. Micronuclei are indicated by white arrows. **(D)** Quantification of cells containing micronuclei from experiments described in **C**. Results are presented as the mean ± SEM from 3 independent experiments. Statistical significance was determined using two-way ANOVA followed by Sidak’s multiple-comparisons test. *P <* 0.01 (**), *P* < 0.001 (***), and *P* < 0.0001 (****). **(E)** HT-29 cells were cocultured with wild-type *pks⁺* (WT) or colibactin-deficient *clbP^S95R^* (Mut) *E. coli* for 4 h at an MOI of 200. Total DNA was immediately isolated and mtDNA integrity was analyzed by long amplicon PCR. mtDNA amplification efficiency was normalized to the amplification efficiency observed following coculture with colibactin-deficient *E. coli*. Results are presented as the mean ± SEM from 4 independent experiments. Statistical significance was determined using an unpaired two-tailed Student’s *t*-test. *P* < 0.001 (***). **(F)** Wild-type and cGAS-knockout HT-29 cells were treated with clb742 for 24 h and then allowed to recover in clb742-free media. At 144 h following clb742 addition, cytoplasmic fractions were isolated and the abundance of the mtDNA D-Loop locus was determined by quantitative PCR. Cytoplasmic mtDNA abundance was normalized to the untreated wild-type (0 h) condition. Results are presented as the mean ± SEM for 3 independent experiments. Statistical significance was determined as in **D**.

mtDNA released into the cytoplasm is a well-established activator of the cGAS-STING pathway (36, 37). We therefore examined whether colibactin exposure damages the mitochondrial genome or promotes cytoplasmic mtDNA accumulation. To assess mtDNA damage, HT-29 cells were cocultured for 4 hours with either *pks+ E. coli* or the colibactin-deficient control strain. Total cellular DNA was then immediately isolated, and the integrity of mtDNA was analyzed by long-amplicon PCR. Because DNA lesions impede DNA polymerase progression, reduced amplification of a long mtDNA fragment provides indirect evidence for the presence of DNA damage. Relative to cells cocultured with the negative control strain, cells cocultured with *pks+ E. coli* showed significantly reduced normalized amplification of the long mtDNA amplicon (Fig. 3E and Fig. S2C), consistent with increased mtDNA damage following colibactin exposure.

We then tested whether mtDNA accumulates in the cytoplasm following colibactin exposure. Wild-type and cGAS-knockout HT-29 cells were treated with clb742 for 24 hours and allowed to recover. Cytoplasmic fractions were then isolated, and their DNA content was analyzed by qPCR using primers targeting two distinct mtDNA loci (Fig. S2D). In wild-type cells, clb742 treatment caused a marked accumulation of cytoplasmic mtDNA during the recovery period (Fig. 3F), whereas total cellular mtDNA levels remained largely unchanged in both wild-type and cGAS-deficient cells (Fig. S2E), suggesting that the increase in cytoplasmic mtDNA reflects redistribution of mtDNA into the cytoplasm rather than an overall increase in mtDNA abundance. Interestingly, clb742-induced cytoplasmic accumulation of mtDNA was largely suppressed in cGAS-knockout cells, suggesting that cGAS activation promotes mtDNA release following colibactin exposure (see below). These results show that colibactin exposure is associated with mitochondrial DNA damage, micronucleus formation, and cytoplasmic accumulation of mtDNA and other dsDNA species.

### Cytoplasmic mtDNA stimulates colibactin-induced cGAS-STING pathway activation

We next asked whether cytoplasmic mtDNA accumulation contributes to cGAS-STING signaling following colibactin exposure. mtDNA was depleted in HT-29 cells using 2’,3’-dideoxycytosine (ddC), a chain-terminating nucleoside analog that preferentially inhibits mtDNA replication by the mitochondrial polymerase Pol γ (38). ddC treatment for 96 hours reduced mtDNA abundance by more than 85%, and mtDNA levels remained low for at least 96 hours after ddC withdrawal (Fig. 4A). Importantly, ddC treatment did not reduce the abundance of the mitochondrial proteins HSP60 and VDAC (Fig. S3A), suggesting that depletion of mtDNA was not accompanied by a substantial reduction in mitochondrial mass. Furthermore, ddC treatment did not impair STING or IRF3 phosphorylation in response to the STING agonist diABZI (Fig. S3B), indicating that mtDNA depletion does not disrupt the ability of STING to activate downstream inflammatory signaling.

**Fig. 4.**
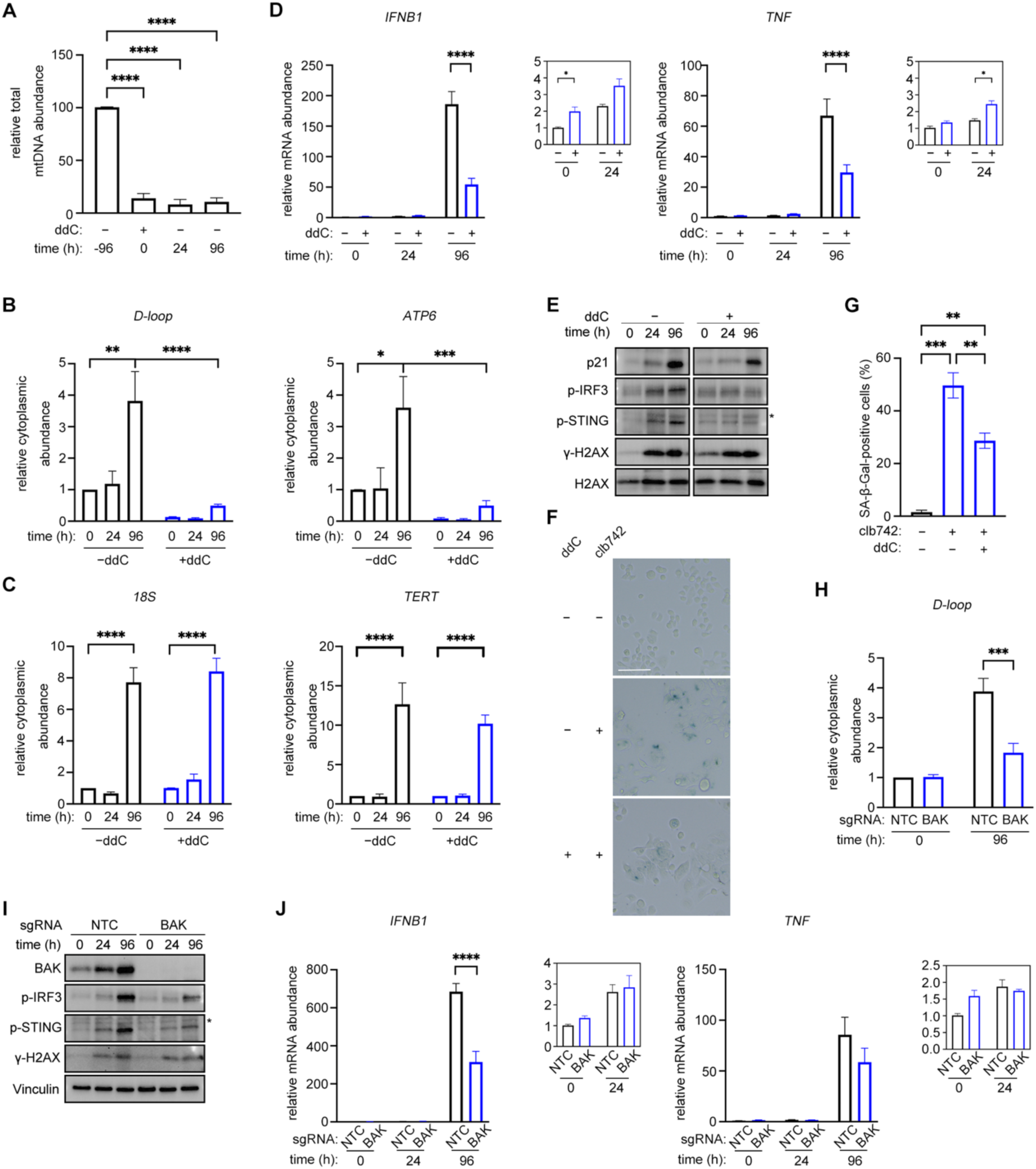
Colibactin-induced mtDNA release promotes cGAS–STING activation and cellular senescence. **(A)** HT-29 cells were treated with ddC for 96 h and then allowed to recover in ddC-free media. Total DNA was isolated and mtDNA D-Loop locus abundance was quantified by quantitative PCR. mtDNA abundance was normalized to untreated cells. Results are presented as mean ± SEM from 3 independent experiments. Statistical significance was determined by one-way ANOVA followed by Dunnett’s multiple-comparisons test. *P* < 0.05 (*)*, P <* 0.01 (**), *P* < 0.001 (***), and *P* < 0.0001 (****). **(B and C)** HT-29 cells were pre-treated with ddC for 96 h, as indicated, and then treated with clb742 for 24 h before being allowed to recover in ddC- and clb742-free media. At the indicated times following clb742 addition, cytoplasmic fractions were isolated and the abundances of the mtDNA D-Loop and *ATP6* loci (B) and the nuclear genome *18S* and *TERT* loci (C) were determined by quantitative PCR. Results are presented as the mean ± SEM from 3 independent experiments. Statistical significance was determined by two-way ANOVA followed by Sidak’s multiple-comparisons test. *P* < 0.05 (*)*, P <* 0.01 (**), *P* < 0.001 (***), and *P* < 0.0001 (****). **(D)** HT-29 cells were treated with ddC and clb742 as in (**B**) and *IFNB1* and *TNF* mRNA expression was determined by quantitative RT-PCR. Relative mRNA expression was normalized to the untreated (-ddC, -clb742; 0 h) condition. Results are presented as the mean ± SEM from 3 independent experiments. Statistical significance was determined as in **B-C**. **(E)** HT-29 cells were treated with ddC and clb742 as in (**B**) and cell lysates were immunoblotted for p21, p-IRF3, p-STING, γH2AX, and H2AX as in Fig. 1C. **(F)** HT-29 cells were treated with ddC and clb742 as in (**B**). 120 h after clb742 addition, cells were stained for SA-β-gal activity. Representative images are shown. Scale bars, 100 µm. **(G)** Quantification of HT-29 SA-β-gal-positive cells from the experiments described in **F**. Results are presented as the mean ± SEM from 3 independent experiments. Statistical significance was determined by one-way ANOVA followed by Tukey’s multiple-comparisons test. *P <* 0.01 (**), *P* < 0.001 (***). **(H)** Non-targeting sgRNA and sgBAK HT-29 knockout cell pools were treated with clb742 for 24 h and then allowed to recover in clb742-free media. At 96 h following clb742 addition, cytoplasmic fractions were isolated and the abundance of the mtDNA D-Loop locus was determined by quantitative PCR. Cytoplasmic mtDNA abundance was normalized to the untreated non-targeting (0 h) condition. Results are presented as the mean ± SEM for 4 independent experiments. Statistical significance was determined as in **B-C**. **(I)** Non-targeting sgRNA and sgBAK HT-29 knockout cell pools were treated with clb742 for 24 h and allowed to recover in clb742-free media. At the indicated times following clb742 addition, cell lysates were immunoblotted for BAK, p-IRF3, p-STING, γH2AX, and vinculin as in Fig. 1C. **(J)** Non-targeting sgRNA and sgBAK HT-29 knockout cell pools were treated with clb742 as in **(I)** and *IFNB1* and *TNF* mRNA expression was determined by quantitative RT-PCR. Relative mRNA expression was normalized to the untreated (0 h) non-targeting sgRNA condition. Results are presented as the mean ± SEM from 3 independent experiments. Statistical significance was determined as in **B-C**.

HT-29 cells were treated with ddC for 96 hours, exposed to clb742 for 24 hours, and then allowed to recover. As expected, ddC treatment reduced clb742-induced cytoplasmic mtDNA accumulation (Fig. 4B). By contrast, ddC treatment did not prevent the cytoplasmic accumulation of DNA derived from the nuclear genome (Fig. 4C). Strikingly, mtDNA depletion reduced clb742-induced phosphorylation of STING and IRF3 and partially suppressed expression of *IFNB1* and *TNF* (Fig. 4D and E). These findings indicate that mtDNA promotes clb742-induced activation of the cGAS-STING pathway. mtDNA depletion also suppressed clb742-induced p21 accumulation and reduced the fraction of SA-β-gal-positive cells (Fig. 4E-G). Together, our results indicate that cytoplasmic mtDNA potentiates colibactin-induced late inflammatory signaling and senescence-associated phenotypes.

Genotoxic stress can promote BAX- and BAK-mediated mitochondrial outer membrane permeabilization (MOMP), which can ultimately release mtDNA into the cytoplasm (39–42). Because clb742 treatment increased BAK abundance in HT-29 cells (Fig. 4H), we tested whether BAK promotes cGAS-STING signaling following colibactin exposure. BAK depletion suppressed clb742-induced cytoplasmic mtDNA accumulation, STING and IRF3 phosphorylation, and *IFNB1* expression (Fig. 4H-J), indicating that BAK promotes activation of the cGAS-STING pathway in cells exposed to colibactin. Overall, our data support a model in which colibactin-induced genotoxic stress activates the cGAS-STING pathway, which promotes BAK-dependent release of mtDNA into the cytoplasm (Fig. 5). Cytoplasmic mtDNA then further stimulates cGAS-STING signaling, enhancing late inflammatory gene expression and senescence-associated phenotypes.

**Fig. 5.**
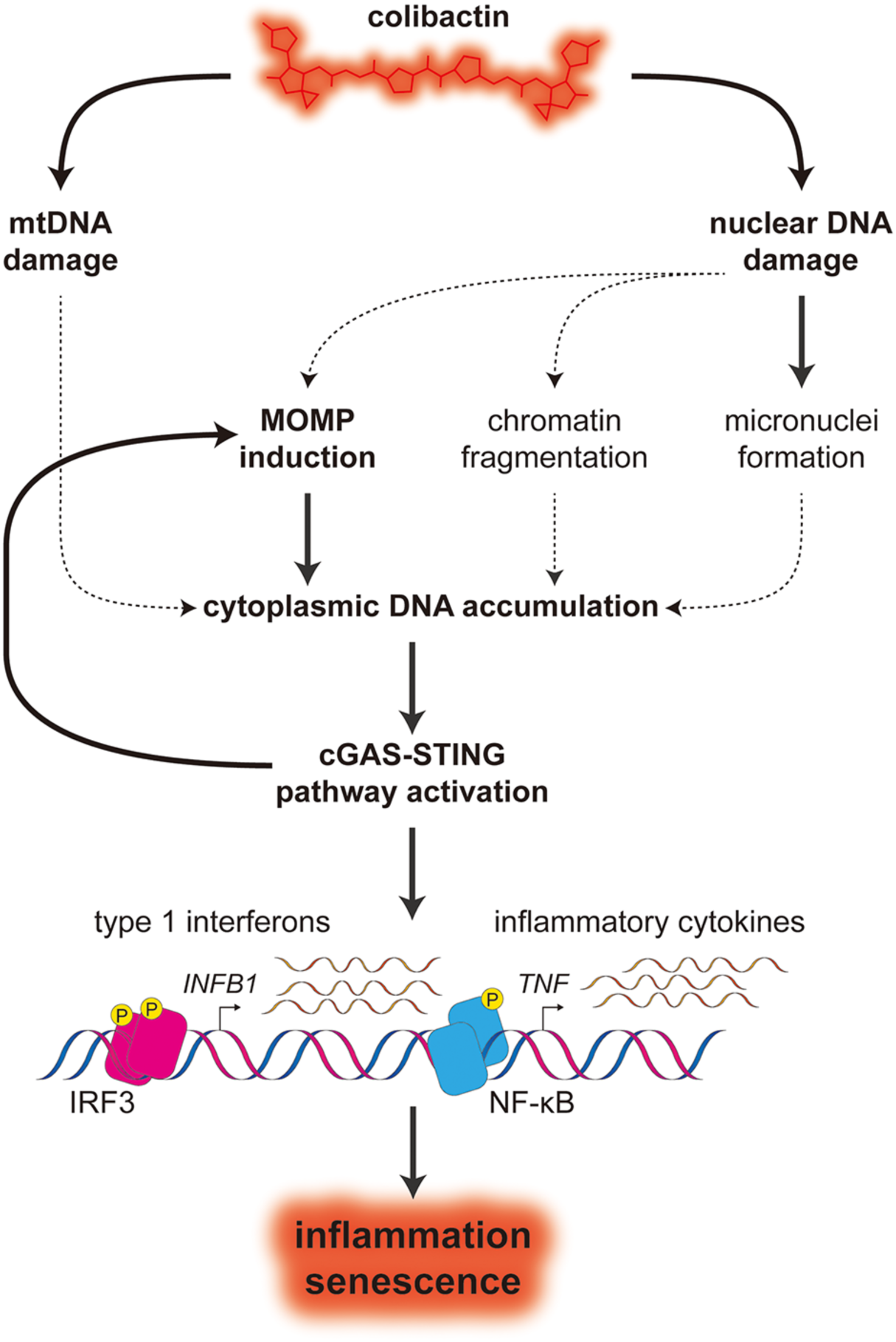
Proposed model linking colibactin-induced genotoxic stress to cGAS–STING-dependent inflammation and senescence. See text for details. Colibactin damages both mitochondrial and nuclear DNA. Nuclear DNA damage results in micronuclei accumulation and MOMP pathway activation, resulting in release of mtDNA into the cytoplasm. Cytoplasmic mtDNA activates the cGAS-STING pathway and downstream induction of inflammatory and senescence programs. Question marks indicate pathways that could contribute to cGAS-STING activation, but have not been experimentally validated.

## Discussion

Colibactin is a potent genotoxin that alkylates DNA and generates mutations associated with CRC. Additionally, colibactin exposure has been linked to inflammatory and senescence-associated phenotypes that may contribute to CRC development and progression. However, the mechanisms connecting colibactin’s genotoxic effects to these responses have remained unclear. Our work establishes a mechanistic link between colibactin-induced DNA damage, inflammatory signaling, and cellular senescence. Consistent with previous work (10, 11), colibactin exposure increased levels of the DNA damage marker γH2AX and promoted micronuclei formation in cultured human cells, consistent with nuclear DNA damage. Colibactin also induced mtDNA damage, although our data do not distinguish whether this damage resulted from direct alkylation of mtDNA or from an indirect mechanism such as increased production of reactive oxygen species. Nuclear and mitochondrial DNA damage was accompanied by cytoplasmic DNA accumulation, increased inflammatory gene expression, and accumulation of senescence-associated markers. Importantly, disruption of cGAS or BAK, as well as depletion of mtDNA, suppressed colibactin-induced inflammatory signaling and senescence-associated phenotypes. Together, these findings indicate that colibactin-induced DNA damage promotes accumulation of cytoplasmic DNA that activates cGAS-STING signaling, with cGAS- and BAK-dependent mtDNA release contributing to the later inflammatory response.

Our study reveals that colibactin-induced host cell responses are coordinated through signaling between the nucleus, mitochondria, and cytoplasm. Following acute colibactin exposure, cells exhibited increased BAK abundance and released mtDNA into the cytoplasm, consistent with engagement of a mitochondrial outer membrane permeabilization pathway. In cells with functional p53, persistent nuclear DNA damage can stabilize and activate p53, which promotes mitochondrial permeabilization through transcriptional induction of BAK and other pro-apoptotic BCL-2 family proteins and through physical interactions with these proteins (39–41, 43). In HT-29 cells, which have defective p53 signaling (44), related genotoxic stress-responsive transcription factors, including p73, could contribute to this response (45, 46). In parallel, cGAS-STING signaling itself may promote mitochondrial permeabilization through phosphorylated IRF3, which can stimulate BAX-dependent MOMP independently of transcription (47–49). Mitochondrial permeabilization would in turn enable release of mtDNA into the cytoplasm, where it can further activate cGAS. Whether colibactin-induced damage to mtDNA enhances cytoplasmic mtDNA accumulation remains to be determined. Thus, our data indicate that colibactin-induced nuclear genotoxic stress initiates inflammatory signaling that promotes mitochondrial permeabilization, establishing a feed-forward loop in which cytoplasmic mtDNA further activates the cGAS-STING pathway.

Notably, colibactin-induced activation of the cGAS-STING pathway occurred in two temporally distinct phases. Within 24 hours of colibactin exposure, we observed phosphorylation of STING and IRF3 accompanied by a modest, 2-to 5-fold induction of *IFNB1* and *TNF*. This early response was not affected by mtDNA depletion or BAK inactivation, implying that it is not driven by mtDNA. Micronuclei began to accumulate during this time frame, raising the possibility that rupture of micronuclei exposes chromatin to cGAS. However, whether micronuclear chromatin is sufficient to activate cGAS remains controversial and may depend on cellular context(35, 50). Alternatively, this early cGAS-STING activation could be driven by nuclear DNA fragments generated during the processing or repair of colibactin-induced lesions. Three to five days after colibactin exposure, cells exhibited dramatic induction of *IFNB1* and *TNF*, exceeding 100-fold in some experiments, along with accumulation of senescence-associated markers. In contrast to the early response, the delayed response coincided with cytoplasmic mtDNA accumulation and was diminished by mtDNA depletion and BAK inactivation, indicating that it is at least partially mediated by released mtDNA. These observations support a biphasic model in which nuclear-derived DNA species promote an early, modest inflammatory response, whereas subsequent mtDNA release contributes to a more robust and persistent inflammatory and senescence-associated program.

A key unresolved question is how DNA repair pathways influence these responses. Colibactin-induced ICLs are repaired by the Fanconi anemia (FA) pathway, and FA pathway-deficient cells are hypersensitive to colibactin (28, 51). Deficits in mismatch repair (MMR) also increase the burden of colibactin-associated mutations (16, 52), suggesting that MMR contributes to the processing or mutagenic consequences of colibactin-induced DNA lesions. By limiting lesion persistence and downstream genome instability, the FA and MMR pathways may constrain colibactin-induced inflammatory signaling. Determining how inactivation of these pathways affects cGAS-STING pathway activation, inflammatory gene expression, and cellular senescence will shed light on how DNA repair capacity shapes host responses to colibactin.

Although our results identify the cGAS-STING pathway as an important mediator of colibactin-induced inflammation and senescence, additional signaling networks likely contribute to these responses. cGAS-knockout cells retained residual induction of *IFNB1* and *TNF* following colibactin exposure and a substantial fraction became positive for SA-β-gal activity, indicating that colibactin alters host-cell physiology through cGAS-independent mechanisms. Persistent DNA damage can trigger cell cycle arrest and apoptosis through p53-dependent pathways that do not require cytoplasmic DNA accumulation (53, 54). DNA damage can also activate NF-κB through the apical DNA damage response kinase ataxia telangiectasia mutated (ATM) and downstream TRAF6- and IKK-dependent signaling (55). Although MAVS depletion did not affect clb742-induced IRF3 phosphorylation, genotoxic stress can engage the RNA sensors RIG-I and MDA5 through increased production of immunostimulatory RNAs derived from endogenous retroelements or mitochondria (56–58), raising the possibility that RNA sensing contributes to the inflammatory response. Colibactin may therefore activate multiple parallel inflammatory pathways, whose relative contributions are likely to depend on cellular and physiological context.

Finally, our study raises the possibility that colibactin-induced cGAS-STING pathway activation influences the tumor microenvironment during colorectal cancer development. cGAS-STING activation can exert opposing tumor-suppressive and tumor-promoting effects depending on cellular context and the duration and intensity of pathway activation (59). Acute signaling can promote tumor-cell senescence and stimulate type I interferon-dependent antitumor immunity. For example, cGAS-STING pathway activation by tumor-derived DNA in dendritic cells can promote activation of tumor-reactive T cells (60). In contrast, persistent cGAS-STING signaling can support tumor progression through chronic NF-κB activation, expression of survival- and metastasis-associated genes, and recruitment of immunosuppressive myeloid cells (61–63). Consistent with a tumor-promoting role for inflammatory signaling in the colon, NF-κB activation enhances tumor formation in a mouse model of colitis-associated CRC (64). Moreover, type I interferons can induce expression of the immune checkpoint pathways that restrain T cell-mediated antitumor responses (65, 66). Together, these observations suggest that the cGAS-STING pathway and other inflammatory signaling pathways influence the consequences of exposure to colibactin-producing bacteria in the gut. Future investigation will be needed to determine whether these pathways promote or restrain colibactin-associated tumorigenesis.

## Methods

### Cell culture, reagents and drug treatments

HT-29, HeLa, and HEK293T cells were cultured in DMEM–GlutaMAX (Gibco, 10569010) containing 4.5 g/L glucose and 1 mM sodium pyruvate, supplemented with 10% fetal bovine serum (FBS; Gibco, A5669801) and 1% penicillin–streptomycin (Gibco, 15140122). Cells were maintained at 37°C in a humidified incubator with 5% CO₂ and were routinely tested to confirm the absence of mycoplasma contamination. Cell numbers and viability were monitored by trypan blue exclusion using a Countess automated cell counter (Thermo Fisher Scientific). HT-29 and HeLa cell lines deficient in cGAS, BAK, or MAVS (cGAS-KO, BAK-KO, and MAVS-KO) were generated by lentiviral delivery of CRISPR/Cas9 constructs followed by puromycin selection, as described below. HT-29 and HeLa cells were treated with the stabilized colibactin analogue clb742 at 20 μM and 10 μM, respectively, for 24 h. clb742 was synthesized as previously described (27). Cells were treated with hydroxyurea (HU; Sigma-Aldrich) at 5 mM for 24 h. Where indicated, cells were treated with the cGAS inhibitor G140 (InvivoGen, inh- g140) at 10 μM beginning 2 h before addition of clb742 treatment. For mitochondrial DNA depletion, cells were treated with 40 μg/mL 2′,3′-dideoxycytidine (ddC; MedChemExpress, HY-17392) for 4 days, and subsequent experiments were performed in ddC-free medium. For STING activation, cells were treated with 0.1 μM diABZI (MedChemExpress, HY-112921B) for 6 h.

### CRISPR–Cas9–mediated gene editing

Single-guide RNAs (sgRNAs) targeting the human cGAS, BAK, and MAVS genes (Table S1) were cloned into the BsmBI site of the LentiCRISPRv2-Puro vector (Addgene #52961). Lentiviral particles were produced by co-transfecting HEK293T cells with LentiCRISPRv2-sgRNA, pRSV-Rev, pMD2.G, and pMDLG/pRRE plasmids using standard calcium phosphate transfection. Viral supernatants were collected 48 h post-transfection, filtered through a 0.45 μm syringe filter, and used to transduce target cells in the presence of 8 μg/mL polybrene (Sigma-Aldrich). Forty-eight hours after infection, cells were selected with puromycin (HT29, 3 μg/mL; HeLa, 2 μg/mL) until all non-transduced control cells were eliminated (approximately 48 h). Single-cell clones were obtained by serial dilution and validated by immunoblotting using the indicated antibodies. Oligonucleotide sequences are listed in Table S1.

### Immunoblotting

Cells were harvested by scraping and lysed by boiling at 98°C for 5 min in 2× SDS loading buffer (100 mM Tris-HCl, pH 6.8, 4% SDS, 20% glycerol, 1% β-mercaptoethanol). Protein lysates were resolved by SDS–PAGE and transferred onto 0.45 μm PVDF membranes. Membranes were blocked with 5% bovine serum albumin (BSA) in TBST (TBS containing 0.1% Tween-20) for 1 h at room temperature and incubated overnight at 4°C with primary antibodies diluted in 1% BSA/TBST. The following antibodies and dilutions were used: anti–phospho-STING (1:2,000; Cell Signaling, D7C3S), anti–phospho-IRF3 (1:1,000; Abcam, ab76493), anti-cGAS (1:4,000; Cell Signaling, D1D3G), anti-γH2AX (1:10,000; Sigma, 05-636), anti-H2AX (1:10,000; Abcam, ab11175), anti-p21 (1:2,000; Proteintech, 10355-1-AP), anti-MAVS (1:1,000; Cell Signaling, 3993), anti-BAK (1:2000, Cell Signaling, 12105) and anti-vinculin (1:5,000; Cell Signaling, E1E9V). After washing, membranes were incubated with HRP-conjugated secondary antibodies (1:10000, Peroxidase AffiniPure® Goat Anti-Rabbit/ Mouse IgG (H+L), Jackson ImmunoResearch) for 1 h at room temperature. Signals were detected using SuperSignal West Dura Extended Duration substrate (Thermo Scientific, 34076) and imaged with a ChemiDoc MP imaging system (Bio-Rad).

### Cell viability assay

Cell viability was assessed using the CellTiter-Glo Luminescent Cell Viability Assay (Promega, G7572) according to the manufacturer’s instructions. Briefly, cells were seeded in 96-well plates in 100 µL media, and an equal volume (100 μL) of CellTiter-Glo reagent was added to each well. Plates were incubated at room temperature for 15 min with gentle agitation, and luminescence was measured using a Tecan GENios microplate reader.

### RNA extraction and qPCR

Total RNA was extracted from HT-29 and HeLa cells using the RNA Isolation Kit (Vazyme, RC112-01) according to the manufacturer’s protocol. One to two micrograms of total RNA were reverse transcribed using the RevertAid First Strand cDNA Synthesis Kit (Thermo Fisher Scientific, K1622). Quantitative PCR was performed using Taq Pro Universal SYBR qPCR Master Mix (Vazyme, Q712-02). Gene expression levels were normalized to *GAPDH*. Each biological replicate was analyzed in technical duplicate. Relative expression was calculated using the 2⁻ΔCt method (67). Primer sequences are listed in Table S1.

### Cell fractionation and cytosolic DNA quantification

Cytosolic DNA was isolated as described previously (68). Briefly, cells were washed with ice-cold PBS and resuspended in digitonin lysis buffer (150 mM NaCl, 50 mM HEPES, pH 7.4, 20 μg/mL digitonin). Samples were rotated at 4°C for 10–20 min to selectively permeabilize the plasma membrane. Lysates were centrifuged at 1,000 × g for 3 min, and the supernatant was transferred to a new tube. This step was repeated twice, followed by centrifugation at 17,000 × g for 10 min to remove residual organelles. Cytosolic DNA was extracted using the QIAquick Nucleotide Removal Kit (QIAGEN, 28306) and quantified by qPCR using mitochondrial- and nuclear-specific primers. Cytosolic mtDNA levels were normalized to total RNA from the same sample. For quantification of total mtDNA, total DNA was extracted using the QIAamp DNA Mini Kit (QIAGEN, 51304), and mtDNA copy number was quantified by qPCR using *D-loop* primers normalized to nuclear *18S* rDNA.

### Immunofluorescence and Microscopy

Cells were grown on glass coverslips and treated as indicated. Cells were fixed with 4% paraformaldehyde for 15 min, permeabilized with 0.5% Triton X-100 for 10 min, and blocked with 3% BSA in PBS. Cytosolic DNA was detected using Pico488 dsDNA Quantification Reagent (Lumiprobe, 42010) diluted a 1x concentration in PBS supplemented with 3% BSA for 30 min. Cells were then washed three times with PBS. Coverslips were mounted with DAPI-containing mounting medium, and images were acquired using Zeiss LSM 800 or LSM 880 confocal microscopes.

### Image analysis

Cells were co-stained with Pico488 and DAPI according to the immunofluorescence protocol. Maximum-intensity projection images were analyzed using ImageJ/Fiji. Nuclear regions of interest (ROIs) were generated from the DAPI channel by threshold-based segmentation followed by binary fill holes and binary open processing. Cytoplasmic ROIs were generated from the Pico488 channel by thresholding and subtraction of the corresponding nuclear ROIs. The integrated density of the Pico488 signal within the cytoplasmic ROI was measured and normalized to the corresponding nuclear Pico488 signal for each cell to determine the relative cytosolic dsDNA level. For micronuclei analysis, DAPI-stained fluorescence images were used to manually count micronuclei-positive cells and total cells in ImageJ/Fiji. The frequency of micronucleus formation was calculated as the percentage of micronucleus-positive cells relative to the total number of cells. For each independent experiment, identical image acquisition settings, including exposure time and gain, were used for both untreated and treated samples.

### Senescence induction and SA-β-Gal staining

For induction of DNA damage–associated senescence, HT-29 and HeLa cells at approximately 40-50% confluence were treated with clb742 for 24 h, followed by replacement with fresh culture medium for an additional 72–120 hours. Senescence-associated β-galactosidase (SA-β-Gal) staining was performed using a commercial kit (Cell Signaling Technology, 9860) according to the manufacturer’s instructions. Images were acquired using an EVOS M3000 imaging system (Invitrogen, Thermo Fisher Scientific).

### Long amplicon mtDNA PCR

Total DNA was extracted from HT-29 cells using the QIAamp DNA Mini Kit (QIAGEN, 51304) according to the manufacturer’s protocol. 15ng of DNA was used as template for PCR by using KAPA Long Range HotStart DNA Polymerase (KAPA Biosystems). Cycle parameters for the long mitochondrial amplification product are as follows: hot start at 75°C for 2 min, initial denaturation at 94°C for 1 min, followed by 26 cycles of denaturation at 94°C for 15 s and extension at 64°C for 12 min. PCR product yield was quantified used Qubit 1X dsDNA HS Assay Kit (Q33230). mtDNA copy number was quantified by qPCR using *D-loop* primers normalized to nuclear *18S* rDNA. Primer sequences are listed in Table S1.

### Cells infection with *Escherichia coli* Strains producing colibactin

Co-culture experiments were performed as previously described (69). The colibactin-producing *Escherichia coli* Nissle 1917 strain (EcN) and its colibactin-deficient derivative (EcN-*clbP^S95R^*) were used for infection. Bacterial cultures were initiated from frozen glycerol stocks and grown overnight. Prior to infection, bacterial density was monitored by measuring the optical density at 600 nm (OD600) (Eppendorf BioSpectrometer# 6135). When the cultures reached an OD600 of approximately 0.5, the bacterial suspension was diluted and added to cells at a multiplicity of infection (MOI) of 100 or 200, as indicated, corresponding to approximately 100 or 200 bacteria per cell. HT-29 and HeLa cells were infected for 4 h, washed thoroughly three times with HBSS, and subsequently cultured in medium supplemented with gentamicin to suppress further bacterial growth. Cells were harvested either 24 hours post-infection to assess early responses to colibactin exposure or 120 hours post-infection to evaluate senescence-associated phenotypes by Western blotting and quantitative PCR.

### Quantification and Statistical analysis

All experiments were performed at least twice independently and yielded consistent results. Statistical analyses were performed using GraphPad Prism. Data are presented as mean ± SEM unless otherwise indicated. For quantitative assays, n represents the number of independent experiments unless otherwise specified. The statistical tests used, sample sizes, and details of multiple-comparison analyses are provided in the corresponding figure legends. A P value < 0.05 was considered statistically significant.

## Disclosure of Delegation to Generative AI

The authors declare the use of generative AI in the research and writing process. According to the GAIDeT taxonomy (2026), the following tasks were delegated to GAI tools under full human supervision: Proofreading and editing

The GAI tool used was: Chat-GPT (GPT 5.6 Sol). Responsibility for the final manuscript lies entirely with the authors. GAI tools are not listed as authors and do not bear responsibility for the final outcomes. Declaration submitted by: Daniel Semlow

## Acknowledgments

We thank members of the Semlow lab, J.L. Camplbell, and E. Bae for helpful discussion and comments on the manuscript. We thank J.P. Nougayrède and É. Oswald for providing the EcN and EcN-*clbP^S95R^*strains. D.R.S. is supported by NIH grant no. R01GM151410, a Shurl and Kay Curci Foundation Research Grant, an Amgen Chem-Bio-Engineering Award, and is a Ronald and JoAnne Willens Scholar and a Vallee Scholar. S.B.H. is supported by NIH grant no. R01CA215553.

## Author Contributions

S.X. performed all experiments under the supervision of D.R.S. T.K. and J.T. generated preliminary data indicating that clb742 activates the cGAS-STING pathway. Y.A., and S.B.H. supplied the clb742 analog. S.X. and D.R.S. wrote the paper with input from all authors.

---

**Supplementary Fig. S1.**
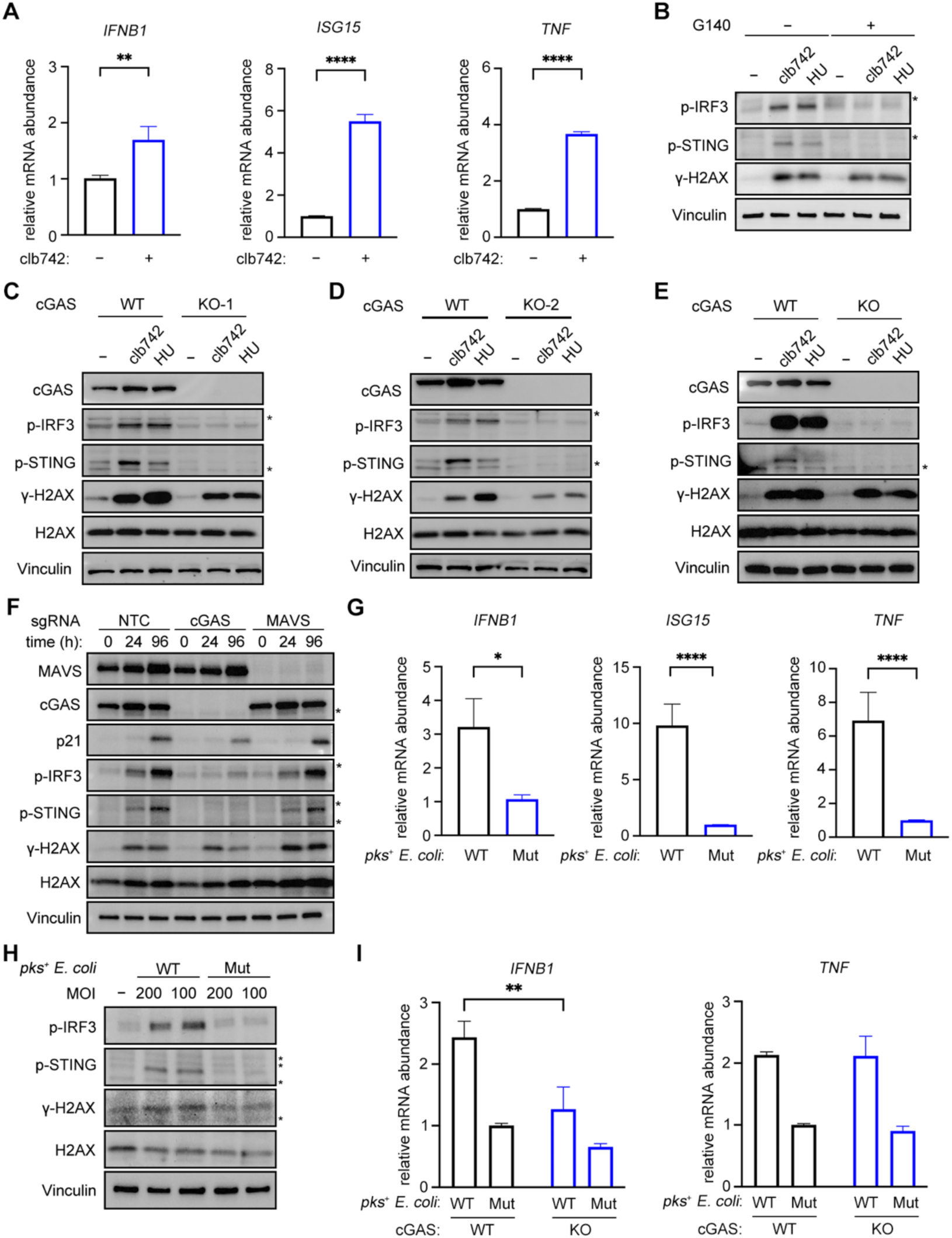
Colibactin-induced cGAS–STING activation in HeLa cells. **(A)** HeLa cells were treated with clb742 for 24 h and *IFNB1*, *ISG15*, and *TNF* mRNA expression was determined by quantitative RT-PCR. Relative mRNA expression was normalized to the untreated condition. Results are presented as the mean ± the standard error of the mean (SEM) from 3 independent experiments. Statistical significance was determined using an unpaired two-tailed Student’s *t*-test. *P* < 0.05 (*)*, P <* 0.01 (**), *P* < 0.001 (***), and *P* < 0.0001 (****). **(B)** HeLa cells were treated with clb742, HU, and G140 for 24 h, as indicated, and cell lysates were immunoblotted for p-IRF3, p-STING, γH2AX, and vinculin. Asterisks indicate non-specific bands. **(C-D)** Wild-type and cGAS-knockout HT-29 single-cell clones (KO-1 and KO-2) were treated with clb742 or HU for 24 h, as indicated, and cell lysates were immunoblotted for cGAS, p-IRF3, p-STING, γH2AX, H2AX, and vinculin. **(E)** Wild-type and cGAS-knockout HeLa cells were treated with clb742 or HU for 24 h, as indicated, and cell lysates were immunoblotted for cGAS, p-IRF3, p-STING, γH2AX, H2AX, and vinculin. **(F)** Non-targeting sgRNA, sgcGAS, and sgMAVS HT-29 knockout cell pools were treated with clb742 for 24 h and allowed to recover in clb742-free media. At the indicated times following clb742 addition, cell lysates were immunoblotted for MAVS, cGAS, p21, p-IRF3, p-STING, γH2AX, H2AX, and vinculin. **(G)** HeLa cells were cocultured with wild-type *pks⁺* (WT) or colibactin-deficient *clbP^S95R^* (Mut) *E. coli* at a multiplicity of infection (MOI) of 200 for 4 h and then allowed to recover in *E. coli*-free media. 24 h following infection, *IFNB1*, *ISG15*, and *TNF* mRNA expression was determined by quantitative RT-PCR. Relative mRNA expression was normalized to the colibactin-deficient *E. coli* condition. Results are presented as the mean ± SEM from 3 independent experiments. Statistical significance was determined as in **A**. **(H)** HeLa cells were cocultured with wild-type *pks⁺* (WT) or colibactin-deficient (Mut) *E. coli* for 4 h at the indicated MOIs. 24 h after infection, cell lysates were immunoblotted for p-IRF3, p-STING, γH2AX, H2AX, and vinculin. **(I)** HT-29 cells were cocultured with wild-type *pks⁺* (WT) or colibactin-deficient *clbP^S95R^* (Mut) *E. coli* at a multiplicity of infection (MOI) of 100 for 4 h and then allowed to recover in *E. coli*-free media. At 24 h following infection, *IFNB1* and *TNF* mRNA expression was determined by quantitative RT-PCR. Relative mRNA expression was normalized to the colibactin-deficient *E. coli* infected wild-type condition. Data are presented as mean ± SEM from 3 independent experiments. Statistical significance was determined using two-way ANOVA followed by Sidak’s multiple-comparisons test, *P <* 0.01 (**).

**Supplementary Fig. S2.**
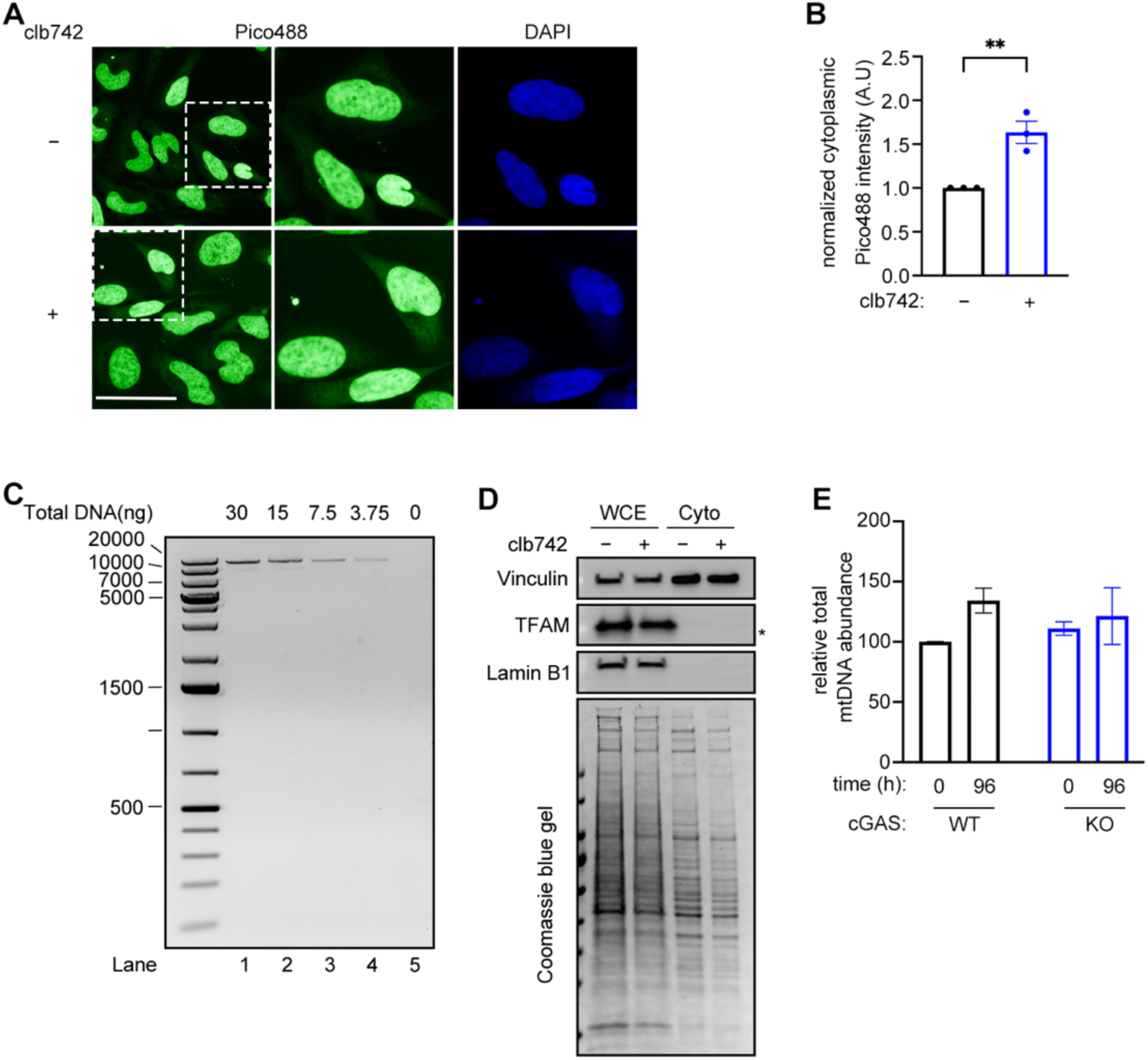
Validation of cytoplasmic DNA accumulation and cytoplasmic fractionation following colibactin treatment. **(A)** HeLa cells were treated with clb742 for 24 h and then stained with Pico488 and DAPI. Regions indicated by boxes are enlarged in the middle and right panels. Representative images are shown. Scale bars, 20 µm. **(B)** Quantification of cytoplasmic Pico488 fluorescence intensity from experiments described in **A**. For each cell, cytoplasmic Pico488 fluorescence intensity was normalized to nuclear Pico488 fluorescence intensity. A total of 152–191 cells were analyzed for the untreated group and 122–134 cells for the clb742-treated group in each independent experiment. Results are presented as the mean ± SEM from 3 independent experiments. Statistical significance was determined using an unpaired two-tailed Student’s *t*-test. *P* < 0.01 (**). **(C)** Long amplicon PCR was performed using the indicated amounts of total HT-29 cell DNA, and amplification products were resolved on an agarose gel and visualized by staining with ethidium bromide. Long amplicon PCR produces a 16.2 kb fragment. **(D)** HT-29 cells were treated with clb742 for 24 h and whole cell lysates and cytoplasmic fractions were separated on an SDS-PAGE gel and immunoblotted for vinculin, TFAM, and Lamin B1, which respectively correspond to markers for the cytoplasm, mitochondria, and nucleus. Total protein was visualized by Coomassie staining. Asterisks indicate non-specific bands. Note the cytoplasmic fractions are enriched for vinculin and depleted for both TFAM and Lamin B1, indicating effective separation of the cytoplasmic fraction from mitochondrial and nuclear fractions. **(E)** Wild-type and cGAS-knockout HT-29 cells were treated with clb742 for 24 h and then allowed to recover in clb742-free media. At 96 h following clb742 addition, total DNA was isolated and the abundance of the mitochondrial D-Loop locus was determined by quantitative PCR. Total mtDNA abundance was normalized to the untreated wild-type (0 h) condition. Results are presented as the mean ± SEM for 3 independent experiments. Statistical significance was determined using two-way ANOVA followed by Sidak’s multiple-comparisons test.

**Supplementary Fig. S3.**
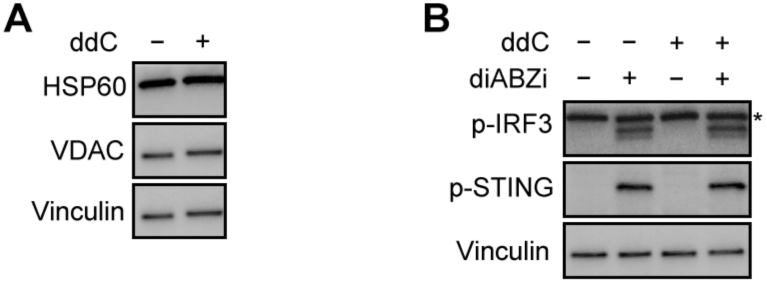
ddC treatment does not alter mitochondrial protein abundance or STING responsiveness. **(A)** HT-29 cells were treated with ddC for 96 h and cell lysates were resolved by SDS-PAGE and immunoblotted for HSP60, VDAC, and vinculin. Note that ddC treatment did not affect the abundance of resident mitochondrial proteins (HSP60 and VDAC), implying that ddC treatment results in mtDNA depletion without affecting overall mitochondrial mass. **(B)** HT-29 cells were pre-treated with ddC for 96 h and then treated with diABZI for 6 h to induce STING activation. Cell lysates were resolved by SDS-PAGE and immunoblotted for p-IRF3, p-STING, and vinculin. Note that ddC treatment did not affect diABZI-induced phosphorylation of IRF3 or STING, implying that mtDNA depletion does not inhibit STING-dependent inflammatory signaling.

**Table S1.**
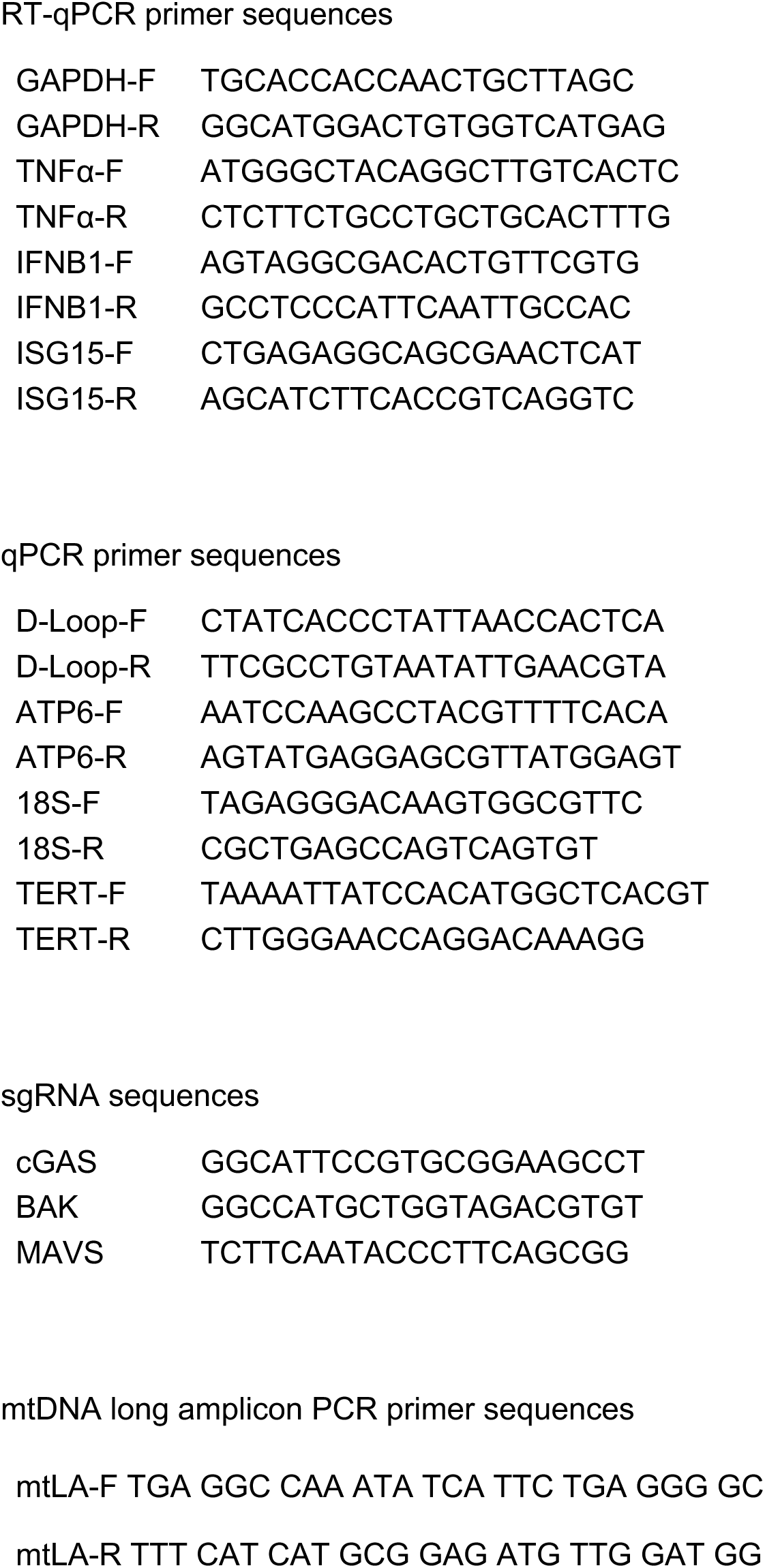
Oligonucleotide sequences used in this study.

